# LKB1 signalling in dendritic cells controls whole-body metabolic homeostasis by limiting T helper 17 priming

**DOI:** 10.1101/2021.10.14.464396

**Authors:** Hendrik J.P. van der Zande, Eline C. Brombacher, Joost M. Lambooij, Leonard R. Pelgrom, Anna Zawistowska-Deniziak, Thiago A. Patente, Graham A. Heieis, Frank Otto, Arifa Ozir-Fazalalikhan, Maria Yazdanbakhsh, Bart Everts, Bruno Guigas

## Abstract

Obesity-associated metaflammation drives the development of insulin resistance and type 2 diabetes, notably through modulating innate and adaptive immune cells in metabolic organs. The nutrient sensor liver kinase B1 (LKB1) has recently been shown to control cellular metabolism and T cell priming functions of dendritic cells (DCs). Here, we report that hepatic DCs from high-fat diet (HFD)-fed obese mice display increased LKB1 phosphorylation and that LKB1 deficiency in DCs (CD11c^ΔLKB1^) worsened HFD-driven hepatic steatosis, systemic insulin resistance and glucose intolerance. Loss of LKB1 in DCs was associated with increased cellular expression of Th17-polarizing cytokines and increased hepatic CD4^+^ IL-17A^+^ Th17 cells in HFD-fed mice. Importantly, IL-17A neutralization rescued metabolic perturbations in HFD-fed CD11c^ΔLKB1^ mice. Mechanistically, disrupted metabolic homeostasis was independent of the canonical LKB1-AMPK axis. Instead, we provide evidence for involvement of the AMPK-related salt-inducible kinase(s) in controlling Th17-polarizing cytokine expression in LKB1-deficient DCs. Altogether, our data reveal a key role for LKB1 signalling in DCs in protection against obesity-induced metabolic dysfunctions by limiting hepatic Th17 differentiation.

## Introduction

Obesity is associated with chronic low-grade inflammation, also known as metaflammation, where continuous overnutrition generates a self-sustained inflammatory loop in metabolic tissues that drives insulin resistance and type 2 diabetes (1). One of the hallmarks of metaflammation is the accumulation of myeloid cells in the main metabolic organs, *i*.*e*. white adipose tissue (WAT), liver and skeletal muscle (2). Macrophage-related cytokines such as tumour necrosis factor (TNF) and interleukin (IL)-1β were shown to inhibit insulin signalling (3, 4) and as such, macrophages are considered key players in the aetiology of tissue-specific insulin resistance. However, dendritic cells (DCs) also accumulate in WAT and liver during obesity and are associated with metabolic dysfunctions. Indeed, depletion of the entire DC population or specific conventional DC (cDC) subsets in different genetic mouse models alleviates adipose tissue and/or hepatic inflammation, although the underlying mechanisms are incompletely understood (5-8).

DCs are specialized antigen presenting cells that govern T cell responses depending on the inflammatory and metabolic microenvironment. Moreover, modulation of T helper cell subsets in metabolic tissues has been shown to play a role in the control of immunometabolic homeostasis. For instance, T helper 2 (Th2) cells and regulatory T cells (Tregs) are enriched in lean, insulin sensitive WAT and contribute to maintenance of tissue-specific insulin sensitivity (9-11). On the contrary, Th17 cells accumulate in WAT and liver during obesity, and are associated with hepatic steatosis and insulin resistance (12-16). In addition, preventing CXCR3-dependent hepatic Th17 accrual and blocking IL-17A signalling using neutralizing antibodies both alleviated non-alcoholic fatty liver disease (NAFLD) (15, 16), suggesting an important contribution of hepatic Th17 cells to NAFLD severity. Although both DCs and T helper cell subsets in metabolic tissues have been associated with control of metabolic homeostasis, little is known about the regulation of DC-mediated T helper cell polarization in these organs during the development of obesity, and its impact on whole-body insulin sensitivity.

DC-mediated priming of Tregs and effector Th1, Th2 and Th17 cells is considered to be driven by metabolic rewiring of DCs in response to environmental cues, which controls co-stimulatory molecule and cytokine expression that shape T helper cell polarization (17). For example, *in vitro* Toll-like receptor (TLR)-activated mature DCs depend on glycolysis for fuelling their anabolic demands, whereas quiescent DCs mainly rely on fatty acid oxidation and mitochondrial oxidative phosphorylation (18). As such, the obesity-induced changes in the metabolic organ microenvironment in which DCs reside may impact their T cell-polarizing capacities and contribute to metaflammation (19).

Among the bioenergetic sensors that regulate DC intrinsic metabolism and function *in vivo*, liver kinase B1 (LKB1) has recently received considerable attention (20-22). The tumour suppressor LKB1 is a serine/threonine kinase that can phosphorylate and activate AMP-activated protein kinase (AMPK) and 12 other members of the AMPK-related family of protein kinases (23, 24), thereby controlling cell growth, survival, polarity and metabolism (25). In DCs, LKB1 was shown to be a critical regulator of effector T cell and Treg priming, thereby maintaining anti-tumour immunity (21, 22). We therefore hypothesized that LKB1 in DCs may connect the changing metabolic microenvironment during obesity to altered T cell priming, thereby impacting whole-body metabolic homeostasis.

In the present study, we investigated the role of LKB1 signalling in DC-mediated T helper cell priming in metabolic tissues and its impact on metabolic homeostasis. We demonstrate that obesity increased LKB1 phosphorylation in hepatic DCs, and that loss of LKB1 in DCs exacerbated metabolic dysfunctions by promoting AMPK-independent Th17 polarization in obese mice. Finally, we identify salt-inducible kinase(s) (SIK) as the possible LKB1 downstream mediator in repressing Th17-polarizing cytokine expression in DCs.

## Methods

### Animals, diet and treatment

All experiments were performed in accordance with the Guide for the Care and Use of Laboratory Animals of the Institute for Laboratory Animal Research and have received approval from the Dutch Central Authority for Scientific Procedures on Animals (CCD; animal license number AVD116002015253). *Itgax*^*cre*^ (CD11c; PMID: 17591855), *Stk11*^*fl/fl*^ (LKB1; PMID: 12226664), *Prkaa1*^*fl/fl*^ (AMPKα1; PMID: 21124450) and WT mice, all on C57Bl/6J background, were purchased from The Jackson Laboratory or Envigo and crossed, housed and bred at the LUMC. Mice were housed in a temperature-controlled room with a 12-hour light-dark cycle and *ad libitum* access to food and tap water under specific pathogen free conditions. To reduce variation due to sex hormone cycles on whole-body metabolism, male mice were used for all *in vivo* experiments. An *a priori* power calculation was done. Analysis was performed blinded to the conditions.

8-16 weeks old age-matched WT, *Stk11*^*fl/fl*^ (CD11c^WT^), *Itgax*^*cre*^ *Stk11*^*fl/fl*^ (CD11c^ΔLKB1^), *Prkaa1*^*fl/fl*^ (CD11c^WT^) and *Itgax*^*cre*^ *Prkaa1*^*fl/fl*^ (CD11c^ΔAMPKα1^) male mice were fed a high fat diet (HFD, 45% energy derived from fat, D12451, Research Diets) for 18-24 weeks as indicated.

For IL-17A neutralization experiments, 12-19 weeks old age-matched CD11c^WT^ and CD11c^ΔLKB1^ mice were systematically randomized over treatment groups based on body weight and fasting blood glucose levels, and fed a HFD for 6 weeks while concomitant biweekly treatment with 200 µg anti-mouse IL-17A (clone 17F3) or IgG1 κ isotype control (clone MOPC-21; both Bio X Cell). At sacrifice, spleen, visceral white adipose tissue (epidydimal; eWAT), brown adipose tissue (intrascapular; BAT) and liver were weighed and collected for further processing.

### Body composition and indirect calorimetry

Body composition was measured by MRI using an EchoMRI (Echo Medical Systems). Indirect calorimetry was performed in groups of 7-8 mice using a Comprehensive Laboratory Animal Monitoring System (Columbus Instruments) with free access to food and tap water. Mice were individually housed at room temperature and a standard 12-hour light/dark cycle was maintained throughout the measurements. Mice were acclimated to the cages for a period of 48 hours before the start of 4 days of measurements at 20 minute intervals. Food intake was assessed by real-time feed weight measurements. Oxygen consumption and carbon dioxide production were measured, and based on this respirometry, energy expenditure (EE), and carbohydrate (CHO) and fatty acid (FA) oxidation were calculated as previously described (26).

### Isolation of leukocytes from spleen

Spleens were collected in 500 μL RPMI 1640 + Glutamax (Life Technolgies), mechanically disrupted, and digested for 20 min at 37 °C in medium supplemented with 1 mg/mL Collagenase D (Roche) and 2000 U/mL DNase I (Sigma-Aldrich). Digested samples were filtered through 100 µm filters and subjected to erythrocyte lysis buffer (0.15 M NH_4_Cl, 1 mM KHCO_3_, 0.1 mM Na_2_EDTA) before counting using a hemocytometer.

### Isolation of stromal vascular fraction from adipose tissue

After a 1 minute transcardial perfusion with PBS post sacrifice, eWAT samples were collected and digested as described previously (27, 28). In short, eWAT samples were minced and incubated for 1 hour at 37°C in an incubator under agitation (60 rpm) in HEPES-buffered Krebs solution, containing 0.5-1 g/L collagenase type I from *Clostridium histolyticum* (Sigma-Aldrich), 2% (w/v) dialyzed bovine serum albumin (BSA, fraction V; Sigma-Aldrich) and 6 mM D-Glucose (Sigma-Aldrich). The samples were passed through a 100 μm filter (Corning Life Sciences) which was washed with PBS supplemented with 2.5 mM EDTA and 5% FCS. After allowing the adipocytes to settle for ∼10 min, the infranatant, consisting of immune cells, was collected and pelleted at 350 x g for 10 min at room temperature. The pellet was treated with erythrocyte lysis buffer, washed with PBS/EDTA/FCS, and counted using a hemocytometer.

### Isolation of leukocytes from liver

Livers were collected and digested as described previously (27, 28). In short, livers were minced and incubated for 45 min at 37°C in RPMI 1640 + Glutamax containing 1 mg/mL collagenase type IV from *Clostridium histolyticum*, 200 U/mL DNase (both Sigma-Aldrich) and 1 mM CaCl_2_. The digested tissues were passed through a 100 μm cell strainer (Corning Life Sciences) which was subsequently washed with PBS/EDTA/FCS. After centrifugation (530 x g, 10 min at 4°C), cells were resuspended in 30 mL PBS/EDTA/FCS and spun down at 50 x g for 3 min at 4°C to pellet the hepatocytes. The supernatant was collected, treated with erythrocyte lysis buffer and CD45^+^ leukocytes were isolated using LS columns and CD45 MicroBeads (35 μL beads per sample, Miltenyi Biotec) according to the manufacturer’s protocol. Isolated liver leukocytes were counted using a hemocytometer.

### Flow cytometry

For assessing LKB1/ACC phosphorylation state in spleen, eWAT and liver, tissues were collected and immediately minced in 1.85% formaldehyde solution (Sigma-Aldrich) and digested as described above. Isolated cell suspensions were permanently permeabilised using 100% methanol for 10 min at -20°C. For other purposes, spleen, eWAT and liver cell suspensions were stained using a Fixable Aqua Dead Cell Stain Kit (Invitrogen) or Zombie UV Fixable Viability Kit (Biolegend) for 20 min at room temperature. Unless sorted or measured alive, cells were fixed for 1 h at 4°C using a FOXP3/Transcription Factor Staining Buffer Set (Invitrogen, for FOXP3 detection) or 15 min at room temperature using a 1.85% formaldehyde solution in PBS (Sigma-Aldrich, for everything else). For detection of intracellular cytokines, isolated cells were cultured for 4 h in RPMI 1640 + Glutamax in the presence of 100 ng/mL phorbol myristate acetate (PMA), 1 µg/mL ionomycin, 10 µg/mL Brefeldin A (all from Sigma-Aldrich). After 4 hours, cells were washed with PBS, stained with Aqua, and fixed as described above. Cell suspensions were first pre-incubated with 2.4G2 antibody (kindly provided by Louis Boon) for blocking Fc receptors and next stained for surface markers in PBS supplemented with 0.5% BSA (Roche) and 2 mM EDTA (Sigma-Aldrich) and antibodies for 30 min at 4°C. For detection of phosphorylated proteins, FOXP3 and intracellular cytokines, cell suspensions were stained in Permeabilization Buffer (eBioscience) instead. Phosphorylated Ser79-ACC and Ser431-LKB1 were stained using unconjugated rabbit-anti-mouse antibodies prior to staining with other antibodies and goat-anti-rabbit-Alexa Fluor 647. Antibody information is provided in Table S1 and gating strategies shown in Figure S1. Cells were measured on a FACSCanto II or LSR II and analysed using FlowJo (Version 10.6, TreeStar).

### Plasma analysis

Blood samples were collected from the tail tip of 4 h-fasted mice using paraoxon-coated glass capillaries. Fasting blood glucose level was determined using a hand-held Glucometer (Accu-Check; Roche Diagnostics) and plasma insulin level was measured using a commercial kit as per manufacturer’s instructions (Chrystal Chem).

### Insulin- and glucose tolerance tests

Whole-body insulin tolerance test (ipITT) and glucose tolerance test (ipGTT) were performed 1 week before sacrifice, as previously described (27, 28). In short, a bolus of insulin (0.75U/kg body mass, NOVORAPID, Novo Nordisk) was administered intraperitoneally (i.p.) to 4 h-fasted mice, after which blood glucose levels were measured at t=0, 15, 30, 45 and 60 min post insulin administration using a Glucometer. For ipGTT, 6 h-fasted mice were injected i.p. with 2g/kg total body mass of D-Glucose (Sigma-Aldrich) and blood glucose was measured at t=0, 20, 40, 60 and 90 min post glucose injection using a Glucometer.

### Histological analysis

Pieces of eWAT and liver (∼30 mg) were fixed in 4% formaldehyde solution (Sigma-Aldrich), paraffin-embedded, sectioned at 4 μm and stained with Hematoxilin and Eosin (H&E). Six fields at 20x magnification (total area 1.68 mm^2^) were used for the analysis of adipocyte size, crown-like structures or hepatic steatosis.

### Hepatic lipid composition

Liver lipids were extracted as previously described (29). Liver triglyceride and total cholesterol concentrations were measured using commercial kits (all from Instruchemie) and expressed as nanomoles per milligram of total protein content using the Bradford protein assay kit (Sigma-Aldrich).

### In vivo DC expansion, isolation and sorting

To expand the DC pool *in vivo*, 2 × 10^6^ Flt3L-secreting B16 melanoma cells (kind gift from Dr. Edward Pearce) in 100 μL HBSS were injected subcutaneously into the flank of mice. After 10 days, spleen, liver and eWAT were harvested, and digested and processed as described earlier. cDC2s were further enriched from single cell suspensions by positive isolation with CD11c Microbeads (Miltenyi Biotec; per manufacturer’s instructions) and FACS sorting (MHCII^+^ CD11c^+^ CD64^-^ F4/80^-^ CD172a^+^ XCR1^-^) on a BD FACSAria using a 100 μm nozzle at 20 PSI. Subsequently, sorted cDC2s were stimulated with 100 ng/mL LPS for 16 h for assessing cytokine expression by RT-qPCR.

### BM-derived DC cultures

Bone marrow-derived DCs were cultured as described previously (22). Briefly, bone marrow cells were flushed from femurs and tibias, and 5 × 10^6^ cells were plated in tissue culture-treated petri dishes (NUNC) in 10 mL of differentiation medium, consisting of RMPI 1640 Glutamax (Gibco) supplemented with 5% FCS (Gibco), 25 nM β-mercaptoethanol (Sigma-Aldrich), 100 U/mL penicillin, 100 μg/mL streptomycin and 20 ng/mL of murine GM-CSF (PeproTech). Medium was refreshed on day 4 and day 7, after which on day 9 non-adherent GMDCs were harvested. 1 × 10^5^ GMDCs were seeded in a round-bottom 96-well plate and rested overnight. The next day, GMDCs were incubated for 2 h at 37°C with 50 µM MARK inhibitor (MARK/Par-1 Activity Inhibitor, 39621; Calbiochem), 50 nM SIK inhibitor (HG-9-91-01; Cayman Chemical), 1 µM NUAK inhibitor (WZ 4003; Tocris) or 5 µM AMPK inhibitor (SBI-0206965; Sigma-Aldrich). After 2 h, LPS and Brefeldin A were added to a final concentration of 100 ng/mL and 10 µg/mL, respectively, and samples were incubated for an additional 4 h at 37°C. After 6 h, cells were stained with a viability kit, fixed using 1.85% formaldehyde and stored at 4°C until further processing for intracellular cytokine detection by flow cytometry.

### RNA-isolation and RT-qPCR

RNA was extracted from LPS-stimulated sorted cDC2s or GMDCs using TriPure RNA Isolation reagent. Total RNA (200-400 ng) was reverse transcribed using the M-MLV Reverse Transcriptase kit (ThermoFisher). Real-time qPCR runs were performed on a CFX96 Real-time C1000 thermal cycler (Biorad) using the GoTaq qPCR Master Mix kit (Promega). Gene expression was normalized to the housekeeping gene *Rplp0* and expressed as fold change compared to CD11c^WT^ samples. A list of primer sequences can be found in Table S2.

### Statistical analysis

All data are presented as mean ± standard error of the mean (SEM). Statistical analysis was performed using GraphPad Prism version 8 for Windows (GraphPad Software) with unpaired t-test, one-way or two-way analysis of variance (ANOVA) followed by Fisher’s post-hoc test. Differences between groups were considered statistically significant at P < 0.05.

## Results

### Obesity induces DC activation in metabolic tissues and increases LKB1 phosphorylation in hepatic DCs

To investigate the role of dendritic cells (DCs) in whole-body metabolic homeostasis during obesity, male C57BL/6J mice were fed a high-fat diet (HFD) for 24 weeks, resulting in significant increases in body weight and fat mass when compared to low-fat diet (LFD)-fed control mice (Fig. 1A-C). Using flow cytometry (Fig. S1), we assessed the frequency and phenotype of DCs in metabolic tissues from lean and obese mice. The number of DCs was found to be significantly increased in WAT but not in the liver from obese mice (Fig. 1D-E). However, DCs from both tissues exhibited increased expression of activation markers (Fig. 1F-G), a feature specific to metabolic tissues as activation status of DCs remained unchanged in the spleen (Fig. S2A). These changes in DC phenotypes were associated with alterations in the T helper cell pool in metabolic tissues. In eWAT, interferon (IFN)γ^+^ Th1 cells were increased at the expense of IL-5^+^ Th2 cells and FOXP3^+^ regulatory T cells (Tregs), while in the liver we detected increased Th1 cells, IL-17A^+^ Th17 cells and Tregs (Fig. 1H-I). In line with unaltered expression of activation markers on splenic DCs, T cell cytokine expression in the spleen was largely unaffected in obese mice (Fig. S2B). These data suggest that the changing microenvironment in metabolic tissues during obesity alters DC activation and, consequently, DC-mediated T cell polarization.

**Figure 1.**
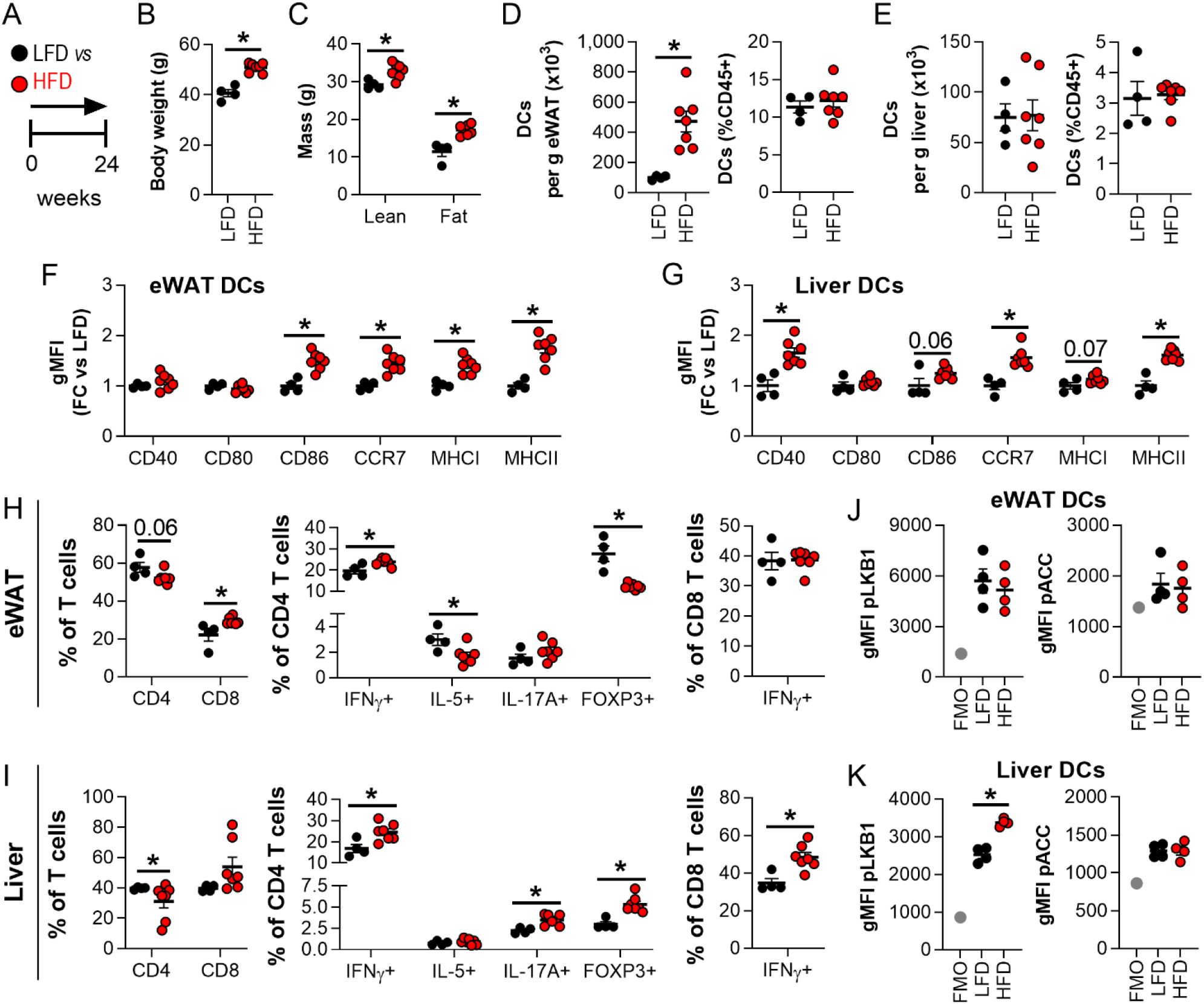
WAT and liver DCs are activated in obese mice. **A**: Mice were fed a low fat diet (LFD; black symbols) or a high fat diet (HFD; red symbols) for 24 weeks. **B-C**: Body weight (*B*) and body composition (*C*) were measured at the end of the experiment. **D-G**: At sacrifice, epidydimal white adipose tissue (eWAT) and liver were collected and immune cells were isolated and analysed by flow cytometry. Absolute numbers of DCs per g tissue and frequencies of total leukocytes in eWAT (*D*) and liver (*E*). Relative expression of indicated DC markers by eWAT (*F*) and liver DCs (*G*). **H-I**: Cells were restimulated with PMA/ionomycin in the presence of Brefeldin A for detection of intracellular cytokines, and were analysed by flow cytometry. CD4 and CD8 T cell, IFNγ^+^ (Th1), IL-5^+^ (Th2), IL-17A^+^ (Th17) and FOXP3^+^ (Treg) CD4 T cell and IFNγ^+^ CD8 T cell percentages in eWAT (*H*) and liver (*I*). **J-K**: eWAT and liver were immediately formaldehyde-fixed after collection and immune cells were isolated. Phosphorylated LKB1 (Ser431) and ACC (Ser79) were measured in DCs from eWAT (*J*) and liver (*K*) by flow cytometry. Full gating strategies are shown in figure S1. Results are expressed as means ± SEM. * P<0.05 vs LFD (n = 4-7 mice per group).

As a bioenergetic sensor, LKB1 was recently shown to be a critical regulator of DC biology and T cell responses *in vivo* (20-22). We next investigated LKB1 signalling in spleen, eWAT and liver DCs by flow cytometry to determine its potential role in tissue-specific DC responses to HFD. Interestingly, we found a marked increase in phosphorylation of Ser431-LKB1 specifically in hepatic DCs from obese mice, suggesting that LKB1 signalling within DC is altered during high-fat feeding, whereas Ser79-ACC phosphorylation, as a proxy for activity of the canonical LKB1 downstream target AMPK, was unchanged (Fig. 1J-K; Fig. S2C). Together, these findings indicate that obesity-induced changes in the hepatic microenvironment may affect LKB1 signalling in DCs which is associated with altered hepatic T cell polarization.

### LKB1 deficiency in DCs aggravates obesity-induced metabolic dysfunctions

To study the role of LKB1 in DCs in the context of obesity-induced metaflammation, we crossed *Stk11*^*flox/flox*^ mice to *Itgax*^*cre*^ mice to generate mice with CD11c-specific deletion of LKB1 as previously described (22). Male conditional knockout (CD11c^ΔLKB1^) and Cre^-^ littermate control (CD11c^WT^) mice were fed an HFD for 18 weeks (Fig. 2A), which did not result in differences in body weight gain or body composition between genotypes (Fig. 2B-E). Food intake, energy expenditure, and carbohydrate (CHO) and fatty acid (FA) oxidation were also not affected by loss of LKB1 in CD11c-expressing cells (Fig. S3). However, despite similar levels at baseline, CD11c^ΔLKB1^ mice developed higher fasting blood glucose levels than CD11c^WT^ littermates after 6 weeks on HFD, which was sustained throughout the experiment (Fig. 2F). Furthermore, whole-body insulin resistance and glucose intolerance were worsened in CD11c^ΔLKB1^ mice, while glucose-induced insulin levels were similar (Fig. 2G-I). Altogether, LKB1 in DCs is important for mitigating insulin resistance and restraining metabolic dysfunctions in mice during HFD-induced obesity.

**Figure 2.**
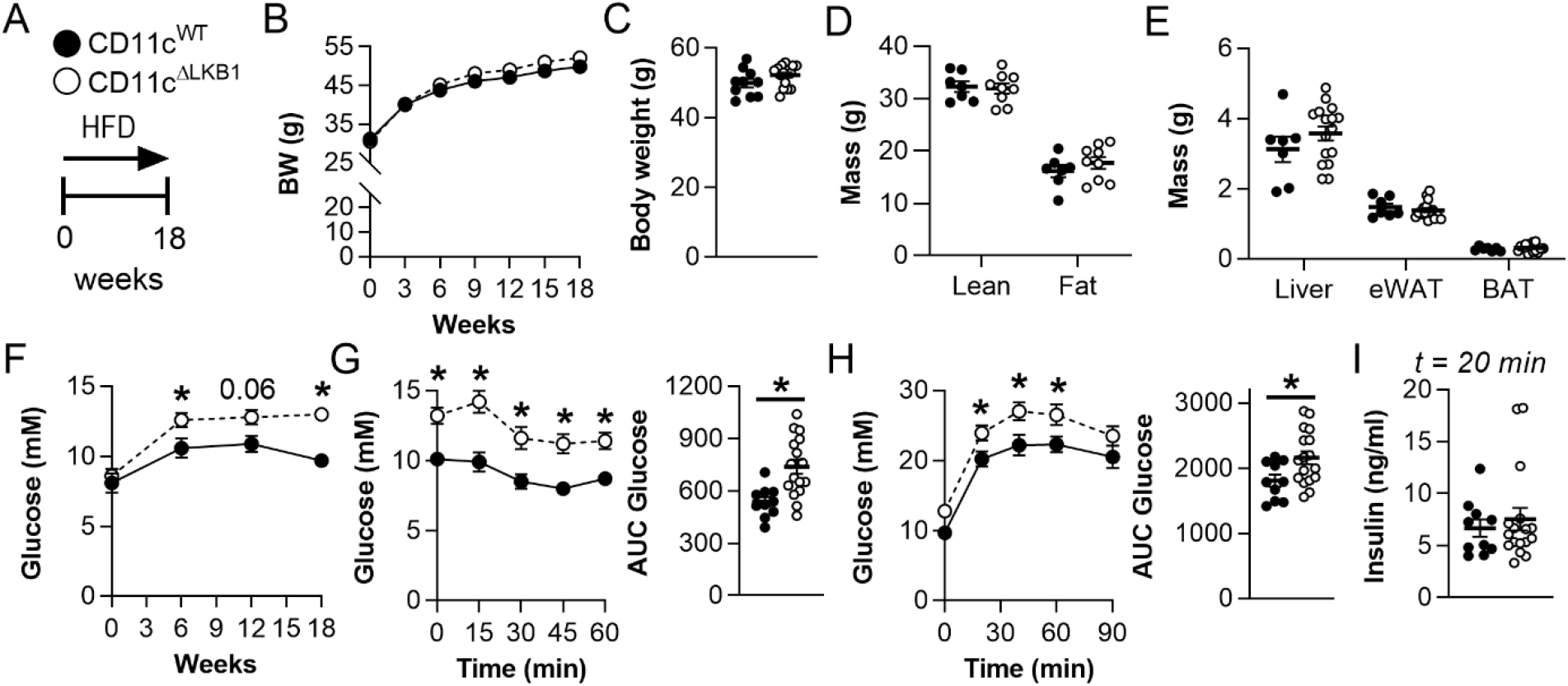
Deletion of LKB1 in DCs aggravates whole-body glucose intolerance and insulin resistance in obese mice. **A**: CD11c^WT^ (black symbols) and CD11c^ΔLKB1^ (open symbols) mice were fed a HFD for 18 weeks. **B-C**: Body weight was monitored throughout the experiment. **D-E**: Body composition (*D*) and weights of liver, eWAT and BAT (*E*) were measured at the end of the experiment. **F**: Fasting blood glucose was measured at the indicated weeks. **G**: An i.p. insulin tolerance test was performed 1 week before sacrifice. Blood glucose levels were measured at the indicated time points and the AUC of the glucose excursion curve was calculated. **H**: An i.p. glucose tolerance test (GTT) was performed 1 week before sacrifice. Blood glucose levels were measured at the indicated time points and the AUC of the glucose excursion curve was calculated. **I**: Plasma insulin was measured at 20 minutes post glucose injection during i.p. GTT. Data shown are a pool of two independent experiments. Results are expressed as means ± SEM. * P<0.05 vs CD11c^WT^ (n = 7-17 mice per group).

### Deletion of LKB1 in DCs promotes hepatic Tregs and Th17 cells and exacerbates hepatic steatosis

We next determined if the exacerbated metabolic dysfunctions observed in obese CD11c^ΔLKB1^ mice could be driven by tissue-specific immunometabolic changes. In eWAT, total leukocyte count and relative abundances of eosinophils, neutrophils, monocytes and CD11c/CD86-expressing macrophages were mostly unaffected (Fig. S4A-F). In line with our previous findings that LKB1-deficient DCs are more migratory (22), we found that the relative abundance of eWAT DCs was decreased in obese CD11c^ΔLKB1^ mice while frequencies of conventional DC (cDC) subsets between genotypes remained similar (Fig. S4G-H). As previous work revealed that LKB1-deficient DCs induced Tregs and effector Th17 cells mostly in lymphoid tissues of lean mice (21, 22), we next assessed whether these T helper subsets are affected in eWAT from obese CD11c^ΔLKB1^ mice. Despite similar CD4 T cell abundance, frequencies of FOXP3^+^ Tregs and IL-17A^+^ Th17 cells within the CD4 T cell pool were increased in eWAT from obese CD11c^ΔLKB1^ mice, while IL-5^+^ Th2 cells were not (Fig. S4I-L). However, when expressed as frequencies of total leukocytes, neither Tregs nor Th17 cells were significantly increased (Fig. S4M-N). Furthermore, adipocyte mean diameter and size distribution were not affected in obese CD11c^ΔLKB1^ mice (Fig. S4O-Q).

In the liver, the abundance of total leukocytes, eosinophils, neutrophils, monocytes and macrophages, in addition to macrophage polarization, was unchanged in obese CD11c^ΔLKB1^ mice when compared to CD11c^WT^ littermates (Fig. 3A; Fig. S5). As was also seen in eWAT, the frequency of DCs was reduced in the livers of CD11c^ΔLKB1^ mice, although the relative abundance of DC subsets remained similar. (Fig. 3B-C). Strikingly, the proportions of liver CD4^+^ Tregs and Th17 cells were significantly increased in mice with LKB1-deficient DCs in comparison to LKB1-sufficient controls (Fig. 3D-I). Moreover, the livers of CD11c^ΔLKB1^ obese mice exhibited enhanced hepatic steatosis when compared to WT littermates (Fig. 3J-K). Consistent with this, hepatic triglyceride (TG) and total cholesterol (TC) levels were also increased (Fig. 3L). Taken together, these results show that deletion of LKB1 in DCs induces a potent increase in Tregs and Th17 cells in the liver and exacerbates hepatic steatosis in obese mice.

**Figure 3.**
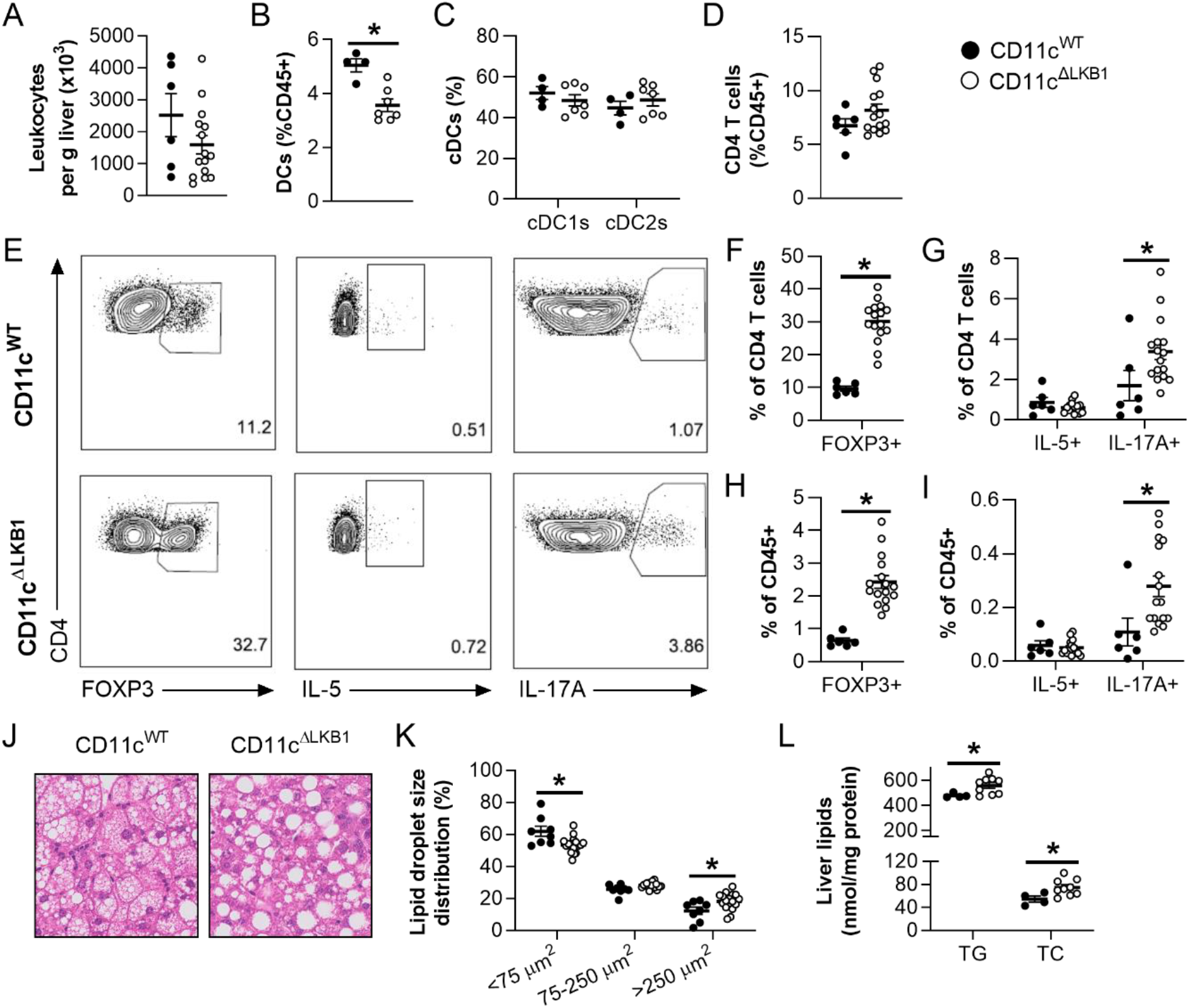
Obese CD11c^ΔLKB1^ mice are more susceptible to HFD-induced hepatic steatosis and have increased hepatic Treg and Th17 cells. CD11c^WT^ (black symbols) and CD11c^ΔLKB1^ (open symbols) mice were fed a HFD for 18 weeks. **A-D**: At sacrifice, liver was collected and immune cells isolated. Total leukocytes per gram liver were quantified (*A*). Percentages of DCs (*B*), cDC subsets (*C*) and CD4 T cells (*D*) were determined by flow cytometry. **E-I**: Liver leukocytes were restimulated with PMA and ionomycin in the presence of Brefeldin A for intracellular cytokine detection. Representative plots (*E*) and percentages of FOXP3^+^ Tregs (*F,H*), IL-5^+^ Th2 and IL-17A^+^ Th17 cells (*G,I*) were determined as frequencies of CD4 T cells (*F,G*) or total leukocytes (*H,I*). **J**: A part of liver was sectioned and H&E stained. **K**: Lipid droplet sizes and size distribution were quantified from H&E-stained slides. **L**: Hepatic triglyceride (TG) and total cholesterol (TC) contents were determined. Data shown are a pool of two independent experiments, except for B, C and L. Results are expressed as means ± SEM. * P<0.05 vs CD11c^WT^ (n = 6-17 mice per group for A, D-K; n = 4-9 mice per group for B, C and L).

### IL-17A neutralization prevents exacerbated obesity-induced metabolic dysfunctions in mice lacking LKB1 in DCs

WAT and liver Th17 cells have consistently been linked to obesity-induced metabolic dysfunctions (13, 30), and hepatic steatosis in particular (12, 14-16). Accordingly, we observed elevated IL-17A-expressing CD4 T cells in the livers of obese mice lacking LKB1 in DCs, a feature that was associated with enhanced hepatic steatosis. Hence, to investigate the contribution of increased Th17 cells to worsened metabolic dysfunctions in CD11c^ΔLKB1^ obese mice, we treated them with either neutralizing antibodies for the Th17 effector cytokine IL-17A or isotype control during the first 6 weeks on HFD (Fig. 4A). IL-17A neutralization did not impact body weight gain (Fig. 4B-C) or hepatic Treg and Th17 cell abundances in CD11c^ΔLKB1^ mice (Fig. S6). However, IL-17A blockade led to significantly improved whole-body insulin sensitivity (Fig. 4D) and reduced hepatic steatosis to comparable levels as CD11c^WT^ littermates (Fig. 4E-G). Thus, increased IL-17A in CD11c^ΔLKB1^ mice plays a central role in promoting liver steatosis and metabolic dysfunctions during HFD-induced obesity. Altogether our findings suggest that LKB1 in DCs mitigates hepatic inflammation during the development of obesity by restraining Th17 priming.

**Figure 4.**
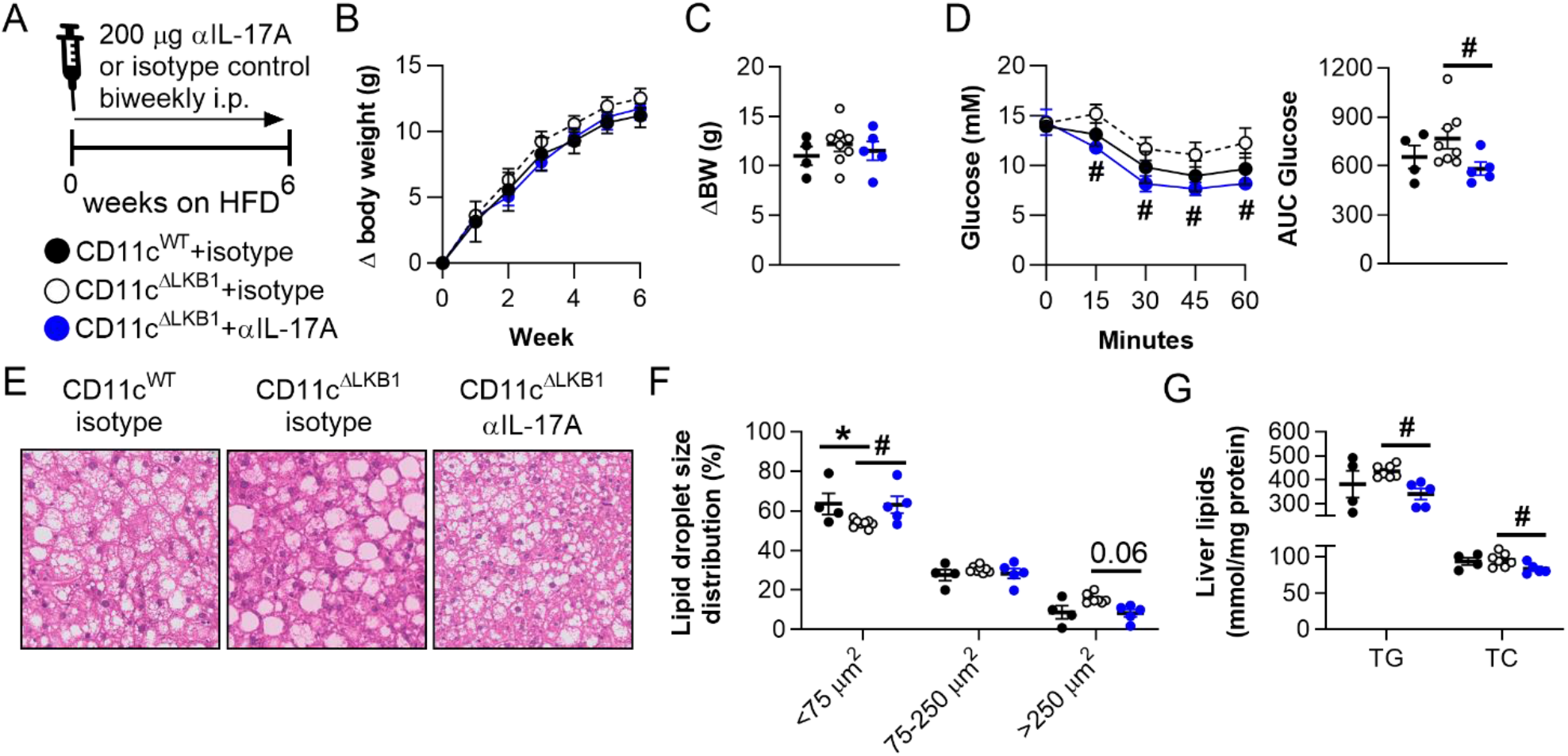
IL-17A neutralization rescued metabolic dysfunctions in CD11c^ΔLKB1^ mice. **A**: CD11c^WT^ (black symbols) and CD11c^ΔLKB1^ mice were fed a HFD for 6 weeks while concomitant biweekly intraperitoneal treatment with IL-17A neutralizing antibodies (blue symbols) or isotype control (open symbols). **B-C**: Body weight gain was monitored throughout the experiment. **D**: An i.p. insulin tolerance test was performed during week 6. **E**: At sacrifice, a piece of liver was sectioned and H&E stained. **F**: Lipid droplet size distribution was quantified from H&E-stained slides. **G**: Hepatic TG and TC content were determined. Data shown are a pool of two independent experiments. Results are expressed as means ± SEM. * P<0.05 vs CD11c^WT^; # P<0.05 vs CD11c^ΔLKB1^ + isotype control (n = 4-8 mice per group).

To explore a direct role for LKB1-deficient DCs in promoting Th17 polarization, we sorted hepatic type 2 conventional DCs (cDC2s), the main CD4 T cell-priming subset shown to induce Th17 priming (31), from lean CD11c^WT^ and CD11c^ΔLKB1^ mice that were subcutaneously injected with Flt3L-secreting B16 melanomas to expand the *in vivo* DC pool (Fig. 5A). Gene expression profiling validated knockout of *Stk11*, encoding LKB1, in hepatic cDC2s from CD11c^ΔLKB1^ mice (Fig. 5C). While surface expression of activation markers on hepatic cDC2s was unchanged during homeostasis (Fig. 5B), LPS-induced expression of the Th17-polarizing cytokines *Il6* and *Il1b* was enhanced in LKB1-deficient hepatic cDC2s when compared to controls, whereas *Il23a* was undetectable and *Tgfb1* unchanged (Fig. 5C). These results show that LKB1 deficiency in DCs promotes production of cytokines, known to favour Th17 polarization, suggesting LKB1 in DCs restrains Th17 polarization by limiting IL-1β and IL-6 production.

**Figure 5.**
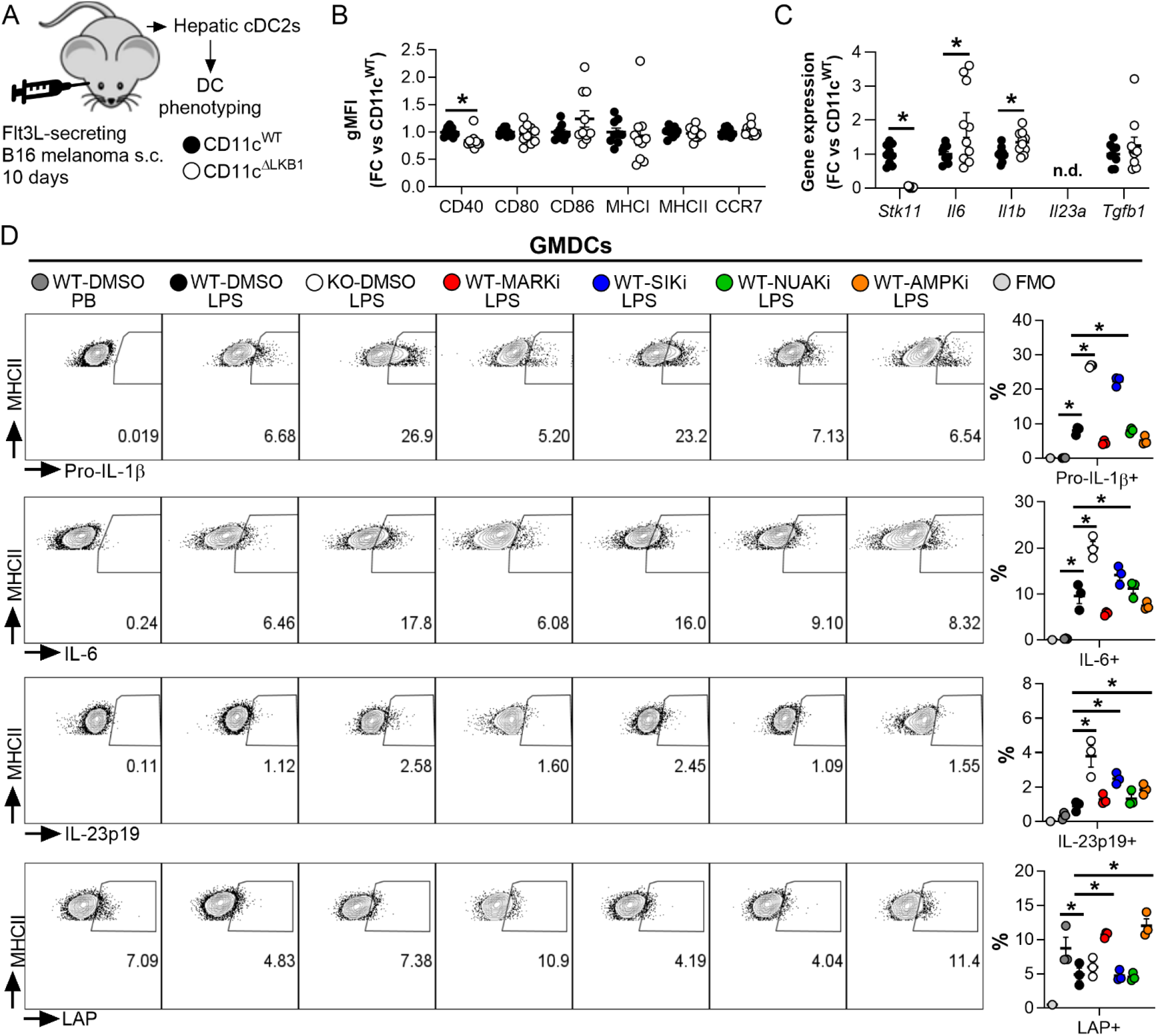
LKB1 deficiency increases Th17-polarizing cytokines expression in DCs, which is mediated through its downstream target SIK. **A-C:** CD11c^WT^ (black symbols) and CD11c^ΔLKB1^ (open symbols) mice were subcutaneously injected with Flt3L-secreting B16 melanomas to expand the DC pool. After 10 days, hepatic cDC2s were FACS sorted for DC phenotyping (*A*). Expression of indicated DC markers was measured by flow cytometry (*B*). Expression of indicated genes was measured by RT-qPCR after *ex vivo* overnight LPS stimulation (*C*). **D:** GM-CSF cultured bone marrow-derived DCs (GMDCs) from CD11c^WT^ (WT) mice were treated with inhibitors targeting LKB1 downstream targets MARKs, SIKs, NUAKs and AMPK for 2 h, before LPS stimulation in the presence of Brefeldin A for 4 h, and compared with CD11c^ΔLKB1^ GMDCs (KO). Pro-IL-1β, IL-6, IL-23p19 and LAP-expressing GMDCs were quantified by intracellular cytokine staining/flow cytometry. Data shown are a pool of three experiments (A-C) or a representative of two experiments (D). Results are expressed as means ± SEM. * P<0.05 vs CD11c^WT^ or as indicated (n = 9-10 mice per group for A-C; n = 3 technical replicates per group for D).

### The LKB1 downstream targets SIKs regulate Th17-polarizing cytokine expression in DCs independent of AMPK

Having demonstrated a role for LKB1 in DCs for preventing excessive obesity-induced metaflammation, we next investigated the signalling mediators downstream of LKB1 responsible for the altered Th17 priming function of DCs. The LKB1-AMPK axis represents a central node in the regulation of cellular energetics, where LKB1 promotes the downstream activation of AMPK though direct phosphorylation of its catalytic α-subunit (25). To assess whether AMPK is involved in the impaired metabolic homeostasis observed in CD11c^ΔLKB1^ obese mice, we generated CD11c^ΔAMPKα1^ mice, in which AMPKα1, the main α-subunit expressed by DCs (32), is deleted in these cells (22). We next fed them and their CD11c^WT^ littermates an HFD for 18 weeks (Fig. S7A). Surprisingly, none of the abovementioned detrimental immunometabolic changes observed in CD11c^ΔLKB1^ obese mice, *i*.*e*. increased fasting glucose levels, glucose intolerance, insulin resistance, and hepatic Tregs and Th17 cells, were recapitulated in HFD-fed CD11c^ΔAMPKα1^ mice (Fig. S7). These data indicate that LKB1-deficiency promotes hepatic Th17 polarization in an AMPK-independent manner.

In addition to AMPK, LKB1 phosphorylates several other downstream AMPK-related kinases including MARK1-4, SIK1-3, NUAK1-2, SNRK and BRSK1-2 (23, 24). We therefore investigated which LKB1 target(s) may contribute to altering DC function by analysing published datasets for their expression in total splenic DCs, as well as mature GM-CSF-elicited bone-marrow DCs (GMDCs). The expression profiles were similar between primary splenic DCs and GMDCs, showing that all these kinases were expressed to a significant level with the notable exception of *Prkaa2* (encoding AMPKα2), confirming that only the catalytic AMPKα1 isoform is expressed by DCs (32), *Mark1* and the members of the BRSK family. Hence, we next determined their role in production of cytokine driving Th17 polarization by DCs. GMDCs were treated with inhibitors of MARK, SIK, NUAK and AMPK families prior to LPS stimulation, and intracellular levels of Th17-polarizing cytokines were assessed by flow cytometry. Largely consistent with liver-derived cDC2s from CD11c^ΔLKB1^ mice, LKB1-deficient GMDCs (Fig. S7C) displayed upregulated LPS-induced expression of pro-IL-1β, IL-6 and IL-23p19 when compared to wild-type GMDCs, whereas latency-associated peptide (LAP) expression, as a proxy for TGF-β production, was unchanged (Fig. 5D). Strikingly, inhibition of SIKs, but not of the other LKB1 downstream kinases, recapitulated the cytokine profile of LKB1-deficient GMDCs (Fig. 5D), identifying SIKs in DCs as potential regulators of Th17 polarization. Collectively, our data indicate that LKB1 signalling in DCs controls hepatic Th17 differentiation and metabolic homeostasis in obese mice, and we propose a role for SIK downstream of LKB1 in repression of Th17-polarizing cytokines.

## Discussion

The bioenergetic sensor LKB1 was recently shown to be a critical regulator of DC metabolism, activation and T cell priming functions (20-22). Whether LKB1 signalling in DCs links the changing immunometabolic microenvironment during obesity with altered DC function and ultimately whole-body metabolic dysfunctions remained unclear. Here, we report that obesity increased LKB1 phosphorylation in hepatic DCs. Deletion of LKB1 from DCs aggravated HFD-induced insulin resistance and hepatic steatosis, and increased hepatic Tregs and Th17 cells in obese mice. These immunometabolic defects were restored by neutralizing the Th17 effector cytokine IL-17A, uncovering a role for LKB1 in restraining DC-mediated pathogenic Th17 cell differentiation, thereby controlling whole-body metabolic homeostasis.

Although DCs accumulate in WAT and liver during obesity and contribute to whole-body insulin resistance (5-8), the underlying mechanisms are incompletely understood. Indeed, obese *Flt3l*^-/-^ mice lacking DCs and *Ccr7*^-/-^ mice with impaired DC migration displayed reduced metaflammation and insulin resistance suggesting they have a central role in the development of metabolic dysfunctions (6, 7). Here, we report that DCs from eWAT and liver, but not spleen, of obese mice display increased expression of activation markers, indicating that the obesity-induced changes in the metabolic tissue microenvironment enhance DC activation. Interestingly, both eWAT and liver DCs from obese mice expressed higher levels of CCR7, suggestive of increased migration to draining lymph nodes where they can prime inflammatory T cells. Consistent with an increased pro-inflammatory activation profile of the DCs, we found that obesity altered the CD4 T helper cell pool in eWAT and liver, but not spleen, favouring Th1 cells at the expense of Th2 cells and Tregs in eWAT, and increasing Th1 cells, Th17 cells and Tregs in the liver. Some of these obesity-induced changes in T helper subsets in metabolic tissues have been reported previously (11, 16). Moreover, XCR1^+^ type 1 conventional DCs (cDC1s), efficient at cross-presenting antigens to CD8 T cells, were reported to increase hepatic steatosis and contribute to liver pathology, which was associated with inflammatory T cell reprogramming in the liver-draining lymph nodes (8). Congruent with this, we found increased HFD-induced IFNγ^+^ CD8 T cells in the liver, most likely resulting from increased cDC1-mediated priming.

In addition to increased DC activation and altered T cell priming in metabolic tissues, we observed a significant increase of Ser431-LKB1 phosphorylation in hepatic DCs of obese mice. LKB1 is phosphorylated at Ser431 by protein kinase C ζ (PKCζ) (33), p90 ribosomal S6 kinase (p90-RSK) and cAMP-dependent protein kinase A (PKA) (34). Although theexact consequence of phosphorylation at this site remains unclear, the residue corresponding to murine Ser431 is conserved in all organisms, suggesting that its phosphorylation may play a role in modulating LKB1 signalling. Despite a lack of a phenotype and normal AMPK activation in knock-in mice carrying a homozygous Ser431 to alanine mutation in LKB1 (35), it has been suggested that Ser431 phosphorylation could promote nuclear export of LKB1 and phosphorylation of some of its cytoplasmic substrates (33, 36).

Interestingly, Ser431-LKB1 phosphorylation was unchanged in splenic and eWAT DCs, indicating that obesity-induced changes in the hepatic microenvironment may specifically alter LKB1 signalling in liver-associated DCs and change its effector functions. Obesity induces persistent changes in the gut microbiota, and endotoxemia through increased gut permeability (37). As a result, the gut and serum metabolome is altered (38, 39), promoting NAFLD pathogenesis through a gut-liver axis (40). Since LPS injection has been reported to acutely increase Ser431-LKB1 phosphorylation in whole lung and liver lysates, and in immortalized Raw264.7 macrophages (41), one may hypothesize that obesity-induced endotoxemia might contribute to increased pLKB1 levels in hepatic DCs from obese mice. Of note, in Raw264.7 macrophages, LPS-induced Ser431-LKB1 phosphorylation suppressed NF-κB signalling, suggesting that increased pLKB1 in hepatic DCs from obese mice may serve as a feedback mechanism to keep inflammation in check. In addition, butyrate, a short-chain fatty acid produced by commensal bacteria that metabolize indigestible fibre, was recently shown to increase Ser431-LKB1 phosphorylation in HepG2 hepatocytes (42), providing conceptual evidence that gut metabolites may alter LKB1 phosphorylation in liver-resident cells. Yet, the underlying mechanisms by which LKB1 signalling is selectively modulated in hepatic DCs during obesity remain to be identified.

Deletion of LKB1 from DCs increased hepatic Tregs and Th17 cells in obese mice. Though predominantly focussed on adipose tissue, the role of Tregs in the regulation of metabolic homeostasis has become controversial over recent years. Seminal work revealed that adipose tissue Tregs are lost during obesity (11, 43), and replenishing the Treg pool through IL-33 treatment or adoptive transfer reduced adipose tissue inflammation and improved metabolic homeostasis (44, 45). However, aging-associated insulin resistance is ameliorated after depletion of adipose tissue Tregs (46), and deletion of the insulin receptor, IL-10 and the transcription factor Blimp1 from Tregs all prevented insulin resistance, in part through promoting adaptive thermogenesis (47, 48). We found a trend for increased adipose tissue Tregs in CD11c^ΔLKB1^ mice, yet eWAT weight, adipocyte size distribution, BAT weight and energy expenditure were unchanged, excluding a role for adaptive thermogenesis in the metabolic phenotype of CD11c^ΔLKB1^ mice. Hepatic Tregs were rather reported to control hepatic inflammation and inhibit NASH development (49). In addition, we clearly demonstrate that neutralizing the Th17 effector cytokine IL-17A rescued the metabolic perturbations in HFD-fed CD11c^ΔLKB1^ mice, indicating an important role for Th17 cells in explaining the metabolic phenotype of obese CD11c^ΔLKB1^ mice.

Indeed, obesity-induced hepatic Th17 cells and IL-17A signalling are consistently reported to impair insulin sensitivity and promote hepatic steatosis (12-16). Th17 differentiation is dependent on the cytokines IL-6, TGFβ, IL-1β and IL-23 (50, 51), but different combinations can lead to different degrees of pathogenicity. In the context of experimental autoimmune encephalitis, IL-6 and TGFβ-induced Th17 cells were not pathogenic, whereas IL-6, IL-1β and IL-23 induced Th17 cells were pathogenic (52, 53). Furthermore, development of Th17 cells *in vivo* is dependent on SIRPα/CD172a expression on DCs (31, 54), a marker of cDC2s that efficiently prime CD4 T cells. We previously showed that LKB1-deficient splenic cDC2s produced higher levels of IL-6 (22). In addition, others showed that mRNA expression of *Il6, Tgfb2* and *Il23a* tended to be increased in LKB1-deficient total splenic DCs compared to WT DCs, whereas *Tgfb1* and *Tgfb3* were decreased or similar (21). Moreover, increased Th17 priming by LKB1-deficient splenic DCs was at least partly dependent on IL-6, but not TGFβ (21). Here, we find that LPS-stimulated LKB1-deficient GMDCs express significantly enhanced levels of IL-6, pro-IL-1β and IL-23p19, while TGFβ production was not affected. In addition, LKB1-deficient hepatic cDC2s express increased levels of *Il6* and *Il1b*, whereas *Tgfb1* was unchanged and *Il23a* undetectable. These data indicate that LKB1-deficient hepatic cDC2s display a cytokine profile that favours the development of pathogenic Th17 cells.

Recent single cell transcriptomics analysis of hepatic Th17 cells from HFD-fed obese mice identified two subsets, of which one was enriched during obesity. The accumulation of this inflammatory hepatic Th17 (ihTh17) subset was regulated through a CXCL9/10-CXCR3 axis, and these cells were sufficient to exacerbate NAFLD pathogenesis through glycolysis-facilitated production of pro-inflammatory cytokines IL-17A, TNF and IFNγ (16). Moreover, increased IL-6, TGFβ, IL-1β and IL-23 levels in steatotic livers were also reported in this study, suggesting involvement of DCs in generating these ihTh17 cells. Given their role in promoting NAFLD pathogenesis, it is tempting to speculate that LKB1-deficient hepatic DCs promote accrual of these ihTh17 cells.

Deletion of AMPKα1 in DCs did not recapitulate the immunometabolic phenotype of CD11c^ΔLKB1^ obese mice. Indeed, we and others have recently shown that LKB1 functions independently of AMPK in governing Tregs and Th17 cell differentiation (21, 22), which corresponds with a growing line of research showing AMPK-independent effects of LKB1 in immune cells (41, 55). We rather show that SIK inhibition, but not of the other DC-expressed AMPK-related kinases, increased expression of IL-6, IL-1β and IL-23 in GMDCs, indicating that an LKB1-SIK axis is likely involved in controlling the expression of cytokines that polarize pathogenic Th17 cells. The SIK family consists of three isoforms, SIK1-3, and is involved in regulating hepatic gluconeogenesis, lipid metabolism and tumorigenesis (56), although its underlying mechanisms are only beginning to be understood. SIKs control the phosphorylation and nucleocytoplasmic transport of class IIa histone deacetylases (HDACs) and cAMP-regulated transcriptional coactivators (CRTCs), identifying a role for SIKs in transcriptional regulation (57). CRTC is a coactivator of cAMP response element-binding protein (CREB) (58), and the promotors of *Il6, Il1b* and *Il23a* all contain CREB binding sites (59-61). It is thus tempting to speculate that inhibition of SIKs may promote CRTC nuclear transport, thereby promoting transcription of Th17-polarizing cytokines. In support of this, SIK1 and SIK3 were shown to control IL-6 production in tumour cells (62), and IL-6 and IL-1β production in Raw264.7 macrophages (63). Conversely, pharmacological inhibition of SIKs was also reported to suppress pro-inflammatory cytokines production in DCs and macrophages (64, 65). As SIK family members display functional redundancy in some settings (57), future studies are required to identify which SIK family member(s) control expression of Th17-polarizing cytokines, and what the mechanistic underpinnings are.

Among the limitations of our study, we cannot formally rule out that deletion of LKB1 in other CD11c-expressing cells, such as macrophages, contributed to the immunometabolic phenotype. Indeed, LysM^cre^-driven LKB1 deletion in macrophages increased LPS-induced pro-inflammatory cytokine production (41). However, these LysM^ΔLKB1^ mice have unchanged Treg numbers (20), indicating that LKB1 deletion from macrophages does not alter T cell polarization *in vivo*. In addition, only partial knockout of *Stk11* was observed in macrophages from CD11c^ΔLKB1^ mice (21, 66), which is likely attributable to lower CD11c expression by macrophages as compared to DCs. Furthermore, we found that expression of the pro-inflammatory/metaflammation-associated markers CD86 and CD11c on both eWAT and liver macrophages was either decreased or unchanged, as was *ex vivo* LPS-induced TNF production by macrophages from CD11c^ΔLKB1^ mice (data not shown). Together, this makes it unlikely that macrophages play a dominant role in the immunometabolic phenotype of CD11c^ΔLKB1^ mice.

Altogether, our data reveal a key role for LKB1 signalling in liver-resident DCs in limiting liver-specific and whole-body metabolic dysfunctions in the context of obesity, by constraining hepatic Th17 accrual. We suggest the involvement of an LKB1-SIK axis to control expression of Th17-polarizing cytokines in DCs, opening interesting therapeutic options in controlling pathogenic Th17 cell development in metaflammation and other hyperinflammatory disorders.

## Acknowledgements

The authors thank Ko Willems van Dijk and Patrick Rensen (Leiden University Medical Center) for allowing the use of the LUMC metabolic phenotyping platform (MRI and metabolic cages). The authors also acknowledge the LUMC Flow cytometry Core Facility (FCF) for technical support and cell sorting assistance.

## Author contributions

HJP van der Zande, EC Brombacher, M. Yazdanbakhsh, B. Everts and B. Guigas conceptualised research; HJP van der Zande, JM Lambooij, F. Otto and B. Guigas analysed data; HJP van der Zande, EC Brombacher, JM Lambooij, LR Pelgrom, A. Zawistowska-Deniziak, TA Patente, G. Heieis, F. Otto and A. Ozir-Fazalalikhan performed research; M. Yazdanbakhsh, B. Everts and B. Guigas supervised the study; and HJP van der Zande, EC Brombacher, B. Everts and B. Guigas wrote the manuscript.

## Figures

**Supplementary Table 1:** Antibodies and reagents for flow cytometry.

| Target | Clone | Conjugate | Source | Identifier |
| --- | --- | --- | --- | --- |
| B220 | RA3-6B2 | FITC | eBioscience | 11-0452 |
| CD3 | 17A2 | APC-eF780 | eBioscience | 47-0032 |
| CD3 | 17A2 | BV605 | Biolegend | 100237 |
| CD3 | 17A2 | FITC | eBioscience | 11-0032 |
| CD4 | GK1.5 | BV650 | BD Biosciences | 563232 |
| CD4 | GK1.5 | PE-Cy7 | eBioscience | 25-0041 |
| CD4 | GK1.5 | PerCP-eFluor 710 | eBioscience | 46-0041 |
| CD8a | 53-6.7 | BV711 | Biolegend | 100759 |
| CD8a | 53-6.7 | PE | eBioscience | 12-0081 |
| CD11b | M1/70 | FITC | eBioscience | 11-0112 |
| CD11b | M1/70 | PE-Cy7 | eBioscience | 25-0112 |
| CD11c | N418 | BV421 | Biolegend | 117330 |
| CD11c | HL3 | FITC | BD Biosciences | 553801 |
| CD11c | HL3 | Horizon V450 | BD Biosciences | 560521 |
| CD11c | N418 | PE-Cy7 | eBioscience | 25-0114 |
| CD19 | MB19-1 | FITC | eBioscience | 11-0191 |
| CD40 | HM40-3 | FITC | eBioscience | 11-0402 |
| CD44 | IM7 | eFluor 450 | eBioscience | 48-0441 |
| CD45 | 30-F11 | BV785 | Biolegend | 103149 |
| CD45.2 | 104 | FITC | Biolegend | 109806 |
| CD45.2 | 104 | eFluor 450 | eBioscience | 48-0454 |
| CD64 | X54-5/7.1 | PE | Biolegend | 139304 |
| CD64 | X54-5/7.1 | PE/Dazzle 594 | Biolegend | 139319 |
| CD64 | X54-5/7.1 | PerCP-Cy5.5 | Biolegend | 139308 |
| CD80 | 16-10A1 | APC | eBioscience | 17-0801 |
| CD86 | GL-1 | APC/Fire 750 | Biolegend | 105045 |
| CD86 | GL-1 | PE | BD Biosciences | 553692 |
| CD172a | P84 | PE | Biolegend | 144011 |
| CD197/CCR7 | 4B12 | PerCP-Cy5.5 | Biolegend | 120116 |
| F4/80 | BM8 | APC | eBioscience | 17-4801 |
| F4/80 | BM8 | BV711 | Biolegend | 123147 |
| FOXP3 | FJK-16s | APC | eBioscience | 17-5773 |
| Goat-anti-Rabbit | Polyclonal | Alexa Fluor 647 | Invitrogen | A21244 |
| GR-1 | RB6-8C5 | FITC | BD Biosciences | 553126 |
| IFNγ | XMG1.2 | FITC | eBioscience | 11-7311 |
| IL-5 | TRFK5 | PE | Biolegend | 504303 |
| IL-6 | MP5-20F3 | APC | Biolegend | 504507 |
| IL-17A | eBio17B7 | PE-Cy7 | eBioscience | 25-7177 |
| IL-17A | TC11-18H10.1 | PerCP-Cy5.5 | Biolegend | 506919 |
| IL-23p19 | fc23cpg | eFluor 660 | Invitrogen | 50-7023 |
| LAP | TW7-16B4 | PerCP-eF710 | Invitrogen | 46-9821 |
| Ly6C | HK1.4 | APC-Cy7 | Biolegend | 128026 |
| MHCI/H-2Kb | AF6-88.5 | Pacific Blue | Biolegend | 116514 |
| MHCII | M5/114 15.2 | Alexa Fluor 700 | Invitrogen | 56-5321 |
| MHCII | M5/114 15.2 | APC-eFluor 780 | eBioscience | 47-5321 |
| MHCII | M5/114 15.2 | FITC | eBioscience | 11-5321 |
| NK1.1 | PK136 | FITC | eBioscience | 11-5941 |
| Phospho-ACC (Ser79) | D7D11 | - | Cell Signaling | 11818S |
| Phospho-LKB1 (Ser431) | C67A3 | - | Cell Signaling | 3482S |
| Pro-IL-1β | NJTEN3 | PE | Invitrogen | 12-7114 |
| Siglec-F | E50-2440 | PE | BD Biosciences | 552126 |
| XCR1 | ZET | BV650 | Biolegend | 148220 |

| Other reagents | Source | Identifier |
| --- | --- | --- |
| LIVE/DEAD™ Fixable Aqua Dead Cell Stain Kit | Invitrogen | L34957 |
| Zombie UV™ Fixable Viability Kit | Biolegend | 423107 |

**Supplementary Table 2:**
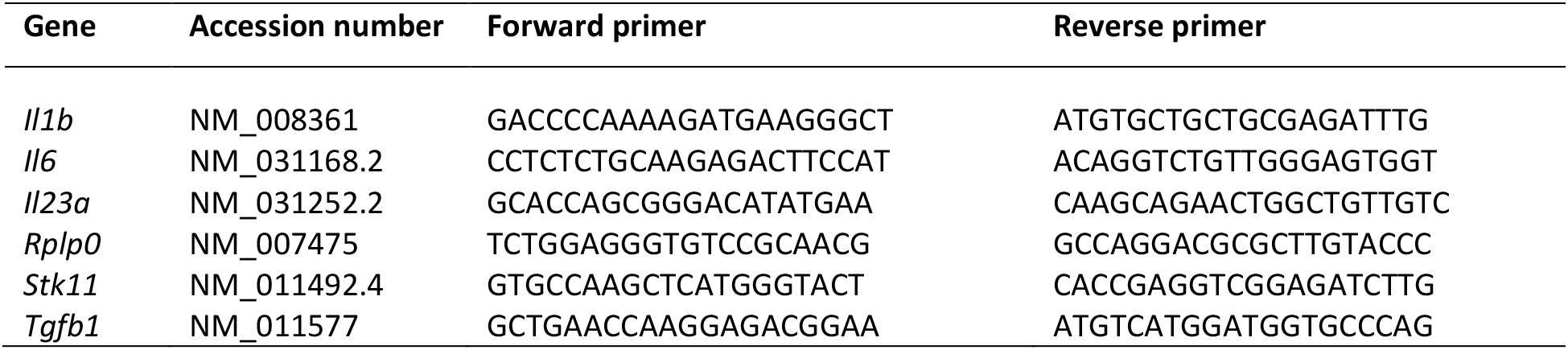
qPCR primers.

## Supplementary figure legends

**Figure S1.**
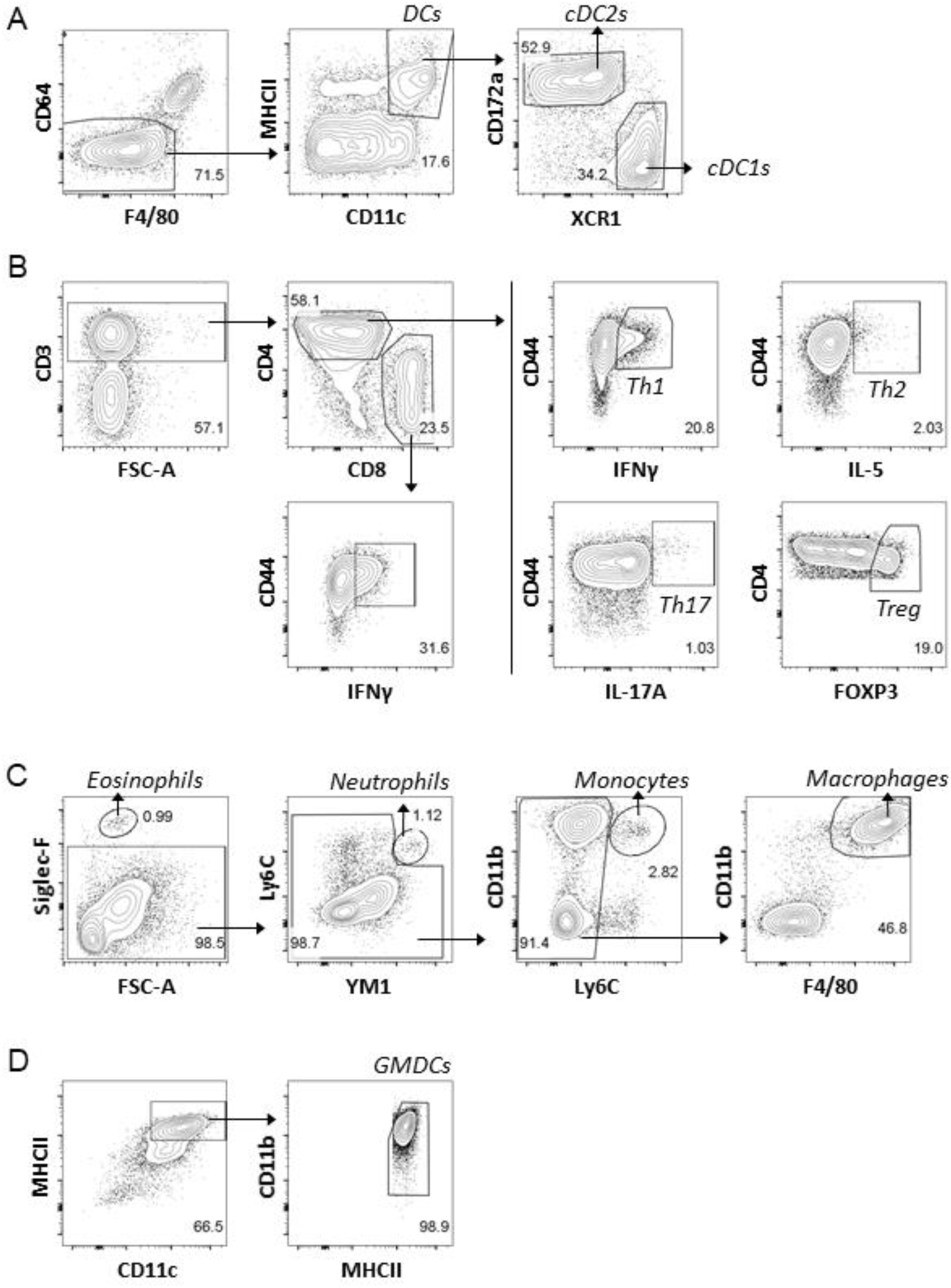
Gating strategies. **A:** Gating strategy for analysis of DCs and cDC subsets. CD11b and CD8a were used as alternatives for CD172a and XCR1, respectively. **B:** Gating strategy for T (helper) cell subsets is shown. **C:** Gating strategy for identification of myeloid cell subsets. **D:** Gating strategy for identification of GMDCs. Isolated cells were pre-gated on live CD45^+^ single cells. For T (helper) cell subset analysis, cells were additionally pre-gated as lineage^-^, which included antibodies directed against B220, CD11b, CD11c, GR-1 and NK1.1. Representative sample was chosen from eWAT samples for A-C. Gating strategies were similar for the indicated cell populations in liver and spleen.

**Figure S2.**
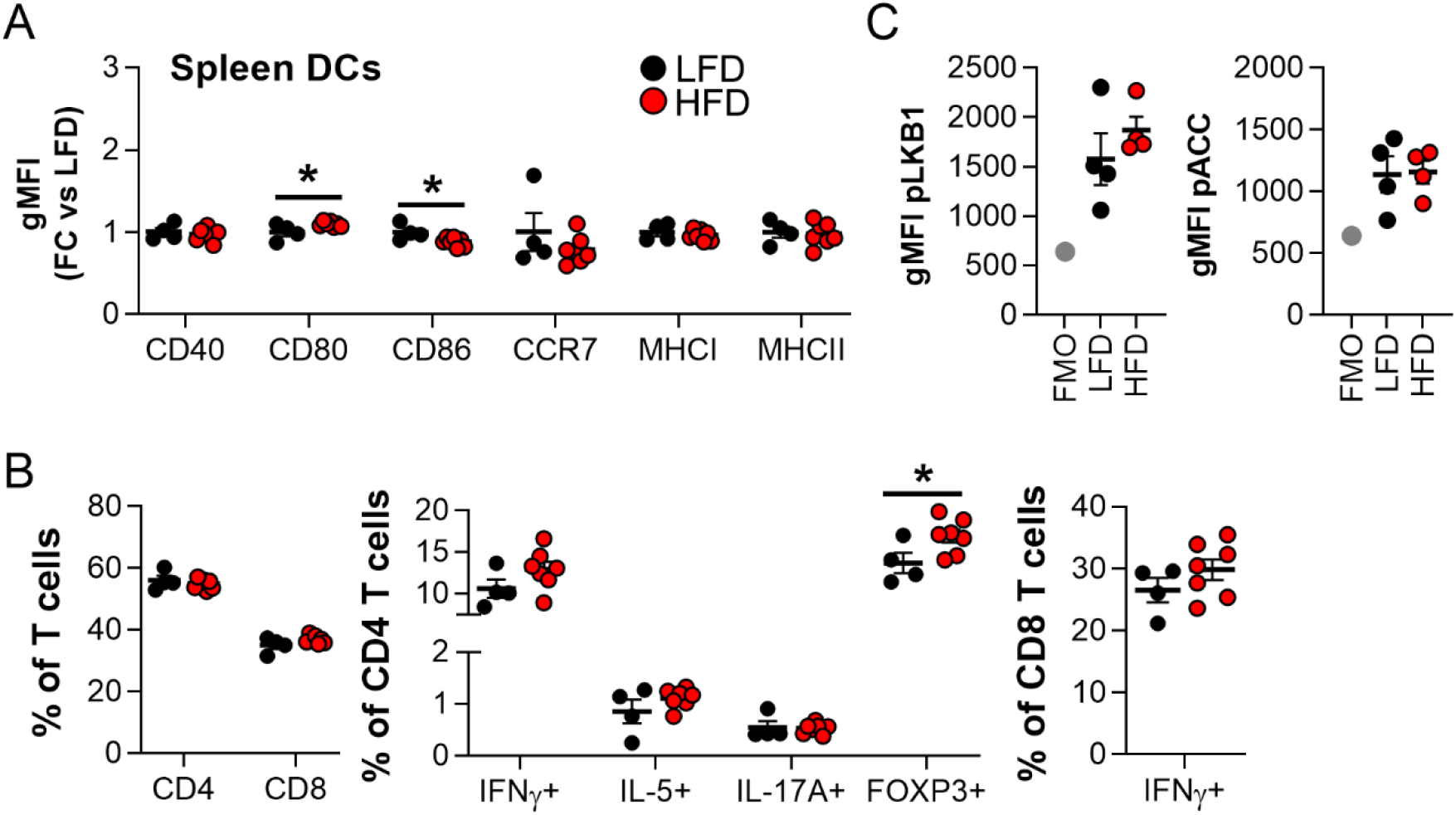
Splenic DCs and T cells are mostly unaffected by obesity. Mice were fed a LFD (black symbols) or a HFD (red symbols) for 24 weeks. **A**: At sacrifice, spleen was collected and immune cells were isolated and analysed by flow cytometry. Relative expression of indicated DC markers by splenic DCs. **B:** Cells were restimulated with PMA/ionomycin in the presence of Brefeldin A for detection of intracellular cytokines, and were analysed by flow cytometry. CD4 and CD8 T cell, Th1, Th2, Th17 and Treg CD4 T cell, and IFNγ^+^ CD8 T cell percentages in spleen. **C**: Spleens were immediately formaldehyde-fixed after collection and immune cells were isolated. Phosphorylated LKB1 (Ser431) and ACC (Ser79) were measured in DCs from spleen by flow cytometry. Results are expressed as means ± SEM. * P<0.05 vs LFD (n = 4-7 mice per group). Related to figure 1.

**Figure S3.**
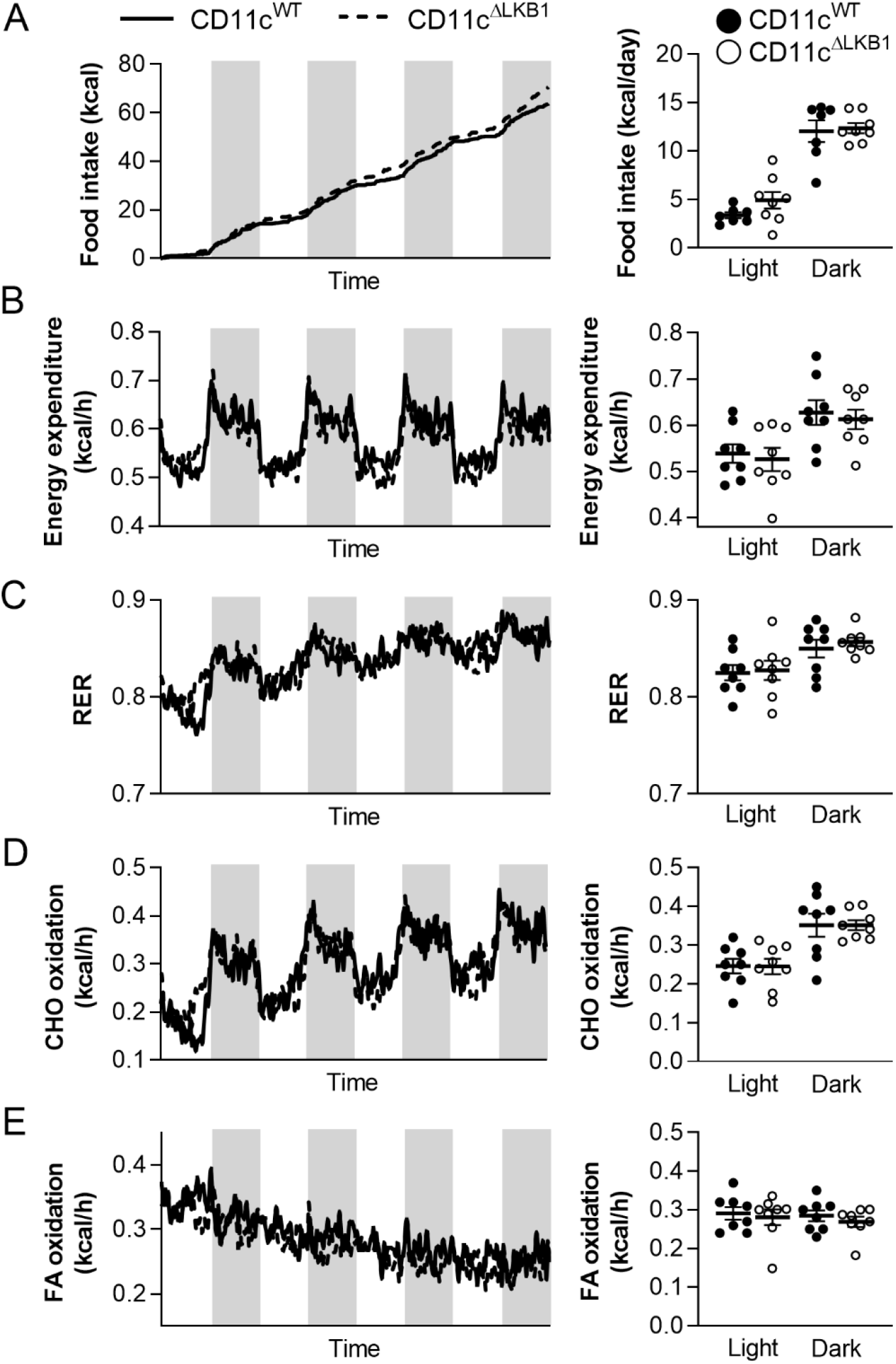
LKB1 deficiency in DCs did not affect food intake and whole-body energy expenditure. CD11c^WT^ (black symbols) and CD11c^ΔLKB1^ (open symbols) mice were fed a HFD for 18 weeks. At week 15, mice were subjected to individual indirect calorimetric measurements using fully automated metabolic cages with free access to food and water. **A-E:** Cumulative food intake (*A*), energy expenditure (EE; *B*), respiratory exchange rate (RER; *C*), carbohydrate (CHO; *D*) and fatty acid (FA; *E*) oxidation were measured for 4 consecutive days (white part = light phase; grey part = dark phase). The daily averages for each of the abovementioned parameters were calculated. Results are expressed as means ± SEM. (n = 7-8 mice per group). Related to figure 2.

**Figure S4.**
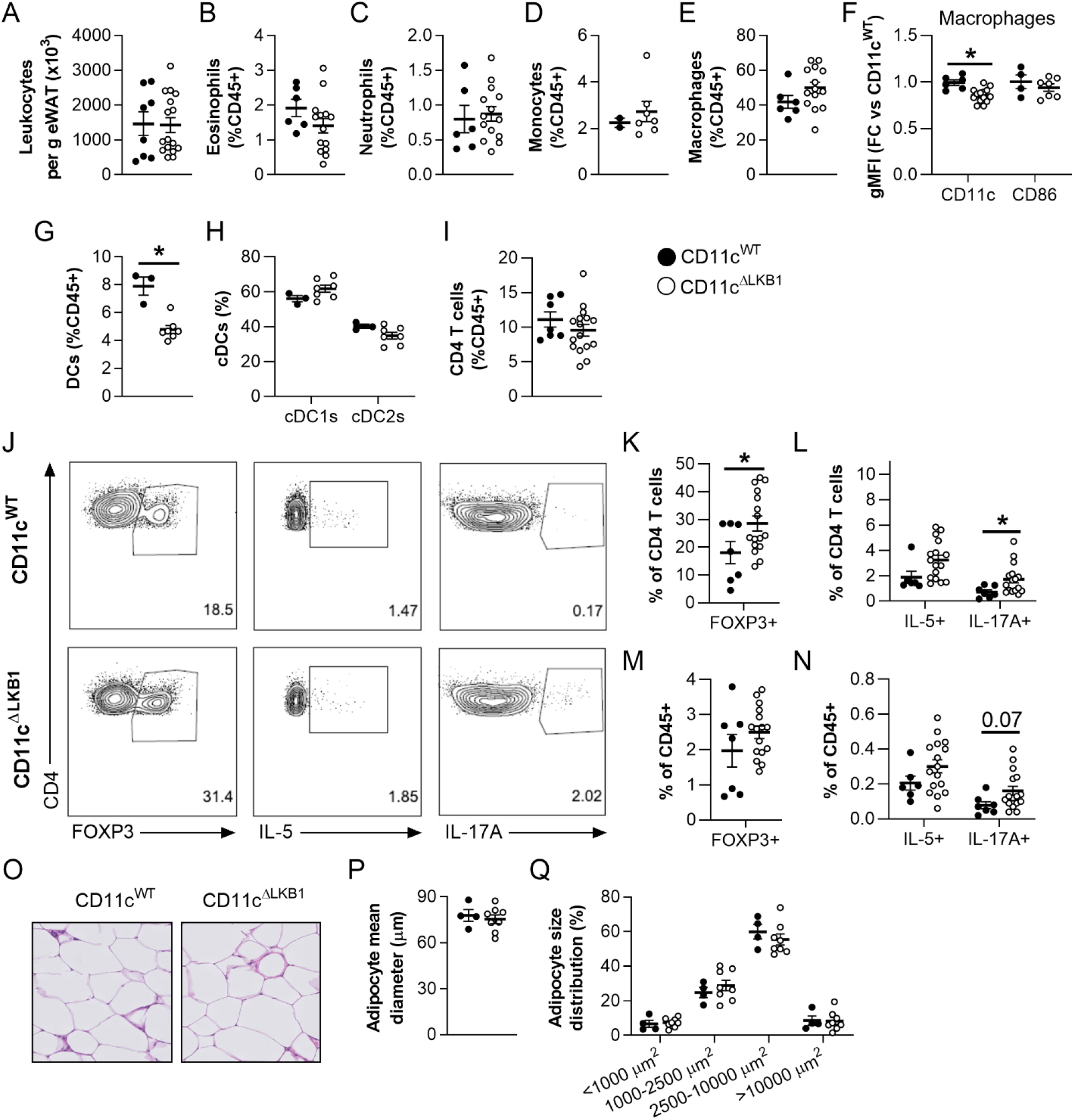
LKB1 deficiency in DCs did not aggravate adipose tissue immunometabolic dysfunctions in obese mice. CD11c^WT^ (black symbols) and CD11c^ΔLKB1^ mice (open symbols) were fed a HFD for 18 weeks. **A-I**: At sacrifice, eWAT was collected and immune cells isolated and analysed by flow cytometry. Total leukocytes per gram eWAT were quantified (*A*). Percentages of eosinophils (*B*), neutrophils (*C*), monocytes (*D*) and macrophages (*E*) in eWAT expressed as frequencies of total leukocytes. Expression of CD11c and CD86 on eWAT macrophages relative to CD11c^WT^ mice (*F*). Abundances of DCs (*G*), cDC subsets (*H*) and CD4 T cells (*I*). **J-N**: eWAT immune cells were restimulated with PMA and ionomycin in the presence of Brefeldin A for intracellular cytokine detection. Representative plots (*J*) and percentages of FOXP3^+^ Treg (*K,M*), IL-5^+^ Th2 and IL-17A^+^ Th17 cells (*L,N*) were determined as frequencies of CD4 T cells (*K,L*) and total leukocytes (*M,N*). **O:** A part of eWAT was sectioned and H&E-stained. **P-Q**: Mean adipocyte diameter (*P*) and adipocyte size distribution (*Q*) were quantified from H&E stained slides. Data shown are a pool of two independent experiments, except for D, F-H and O-Q. Results are expressed as means ± SEM. * P<0.05 vs CD11c^WT^ (n = 7-16 mice per group for A-C, E and I-N; n = 3-8 mice per group for D, F-H and O-Q).

**Figure S5.**
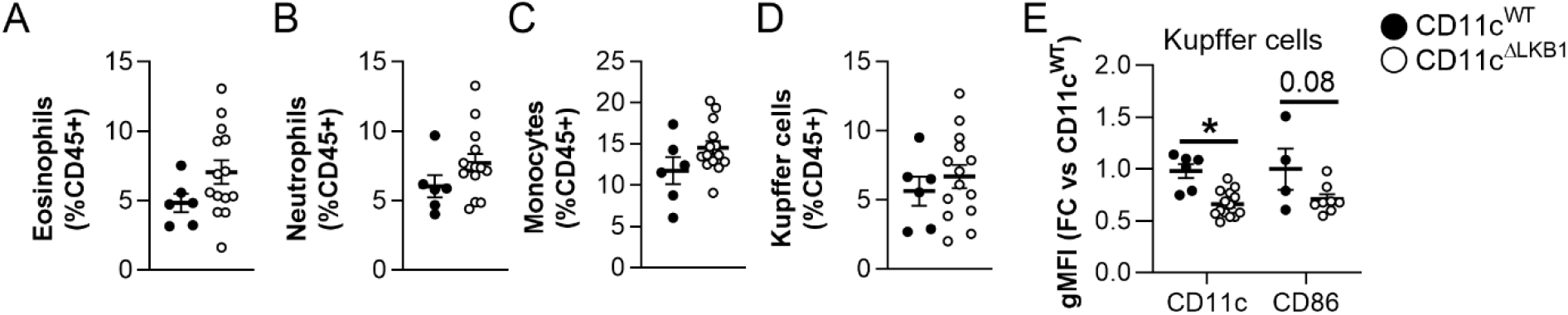
Effects of LKB1 deletion from DCs on myeloid cell subsets in the liver. CD11c^WT^ (black symbols) and CD11c^ΔLKB1^ (open symbols) mice were fed a HFD for 18 weeks. At sacrifice, liver was collected and immune cells were isolated and analysed by flow cytometry. **A-E**: Percentages of hepatic eosinophils (*A*), neutrophils (*B*), monocytes (*C*) and Kupffer cells (*D*) expressed as frequencies of total leukocytes. Expression of CD11c and CD86 on Kupffer cells, expressed as fold change *vs* CD11c^WT^ (*E*). Data shown are a pool of two independent experiments, except for CD86 expression in E. Results are expressed as means ± SEM. * P<0.05 vs CD11c^WT^ (n = 4-14 mice per group). Related to figure 3.

**Figure S6.**
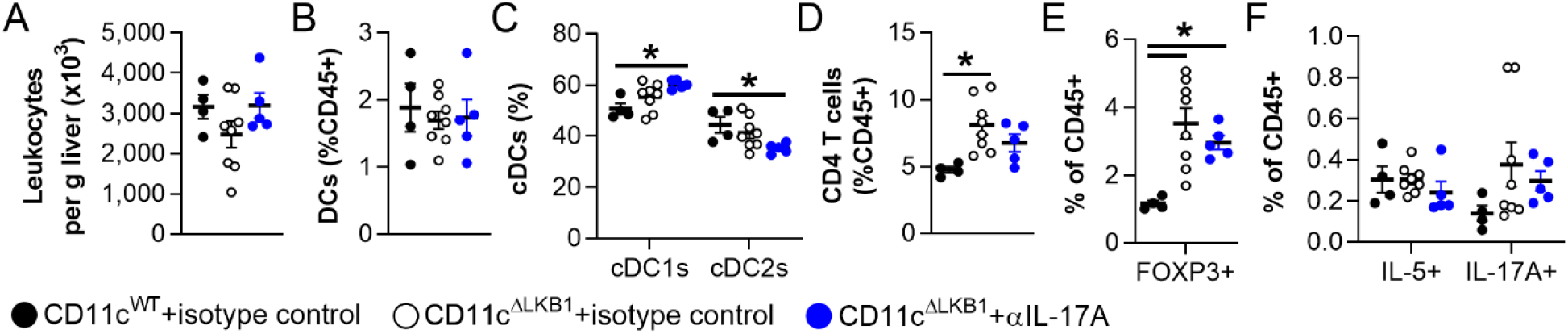
Effects of IL-17A neutralization on hepatic immune cells. Mice were treated as described in the legend of figure 5. **A-D**: At sacrifice, liver was collected and immune cells were isolated and phenotyped by flow cytometry. Total number of leukocytes per gram liver (*A*), and frequencies of DCs (*B*), cDC subsets (*C*) and CD4 T cells (*D*) were determined. **E-F**: Hepatic leukocytes were restimulated with PMA/ionomycin in the presence of Brefeldin A for detection of intracellular cytokines. Abundance of FOXP3^+^ Tregs (*E*), IL-5^+^ Th2 cells and IL-17A^+^ Th17 cells (*F*) were determined as frequencies of total leukocytes. Data shown are a pool of two independent experiments. Results are expressed as means ± SEM. * P<0.05 vs CD11c^WT^ (n = 4-8 mice per group). Related to figure 4.

**Figure S7.**
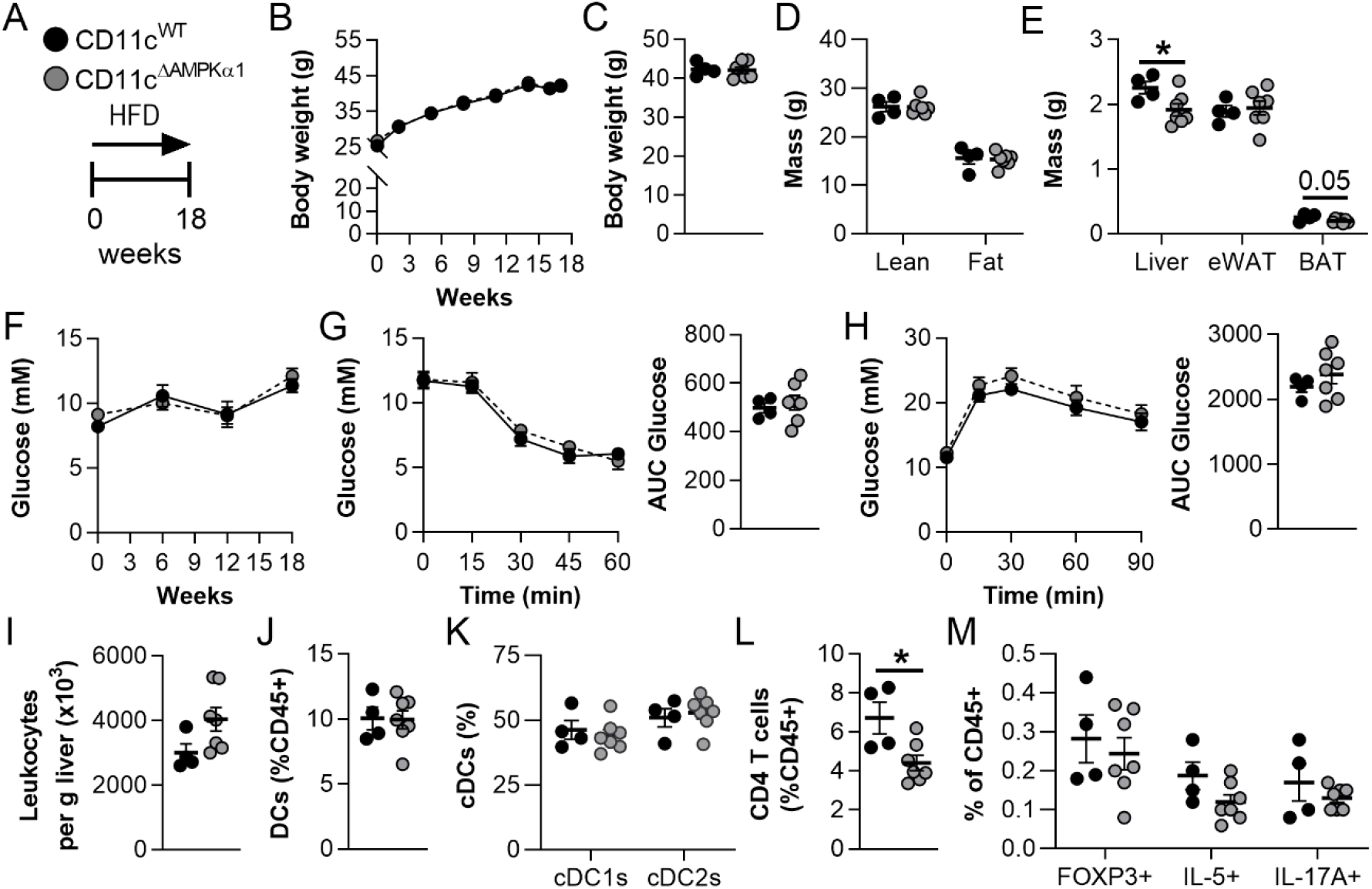
Deletion of AMPKα1 from DCs did not recapitulate the immunometabolic phenotype of CD11c^ΔLKB1^ mice. **A**: CD11c^WT^ (black symbols) and CD11c^ΔAMPKα1^ mice (grey symbols) were fed a HFD for 18 weeks. **B-C**: Body weight was monitored throughout the experiment. **D-E**: Body composition (*D*) and weights of liver, eWAT and BAT (*E*) were measured at the end of the experiment. **F**: Fasting blood glucose was measured at the indicated weeks. **G**: An i.p. insulin tolerance test was performed 1 week before sacrifice and AUC calculated. **H**: An i.p. glucose tolerance test was performed 1 week before sacrifice and AUC calculated. **I-L**: At sacrifice, liver was collected and immune cells isolated. Total leukocytes per gram liver were quantified (*I*). Percentages of DCs (*J*), cDC subsets (*K*) and CD4 T cells (*L*) were determined by flow cytometry. **M**: Liver leukocytes were restimulated with PMA and ionomycin in the presence of Brefeldin A for intracellular cytokine detection. Percentages of FOXP3^+^ Treg, IL-5^+^ Th2 and IL-17A^+^ Th17 cells were determined as frequencies of total leukocytes. Results are expressed as means ± SEM. * P<0.05 vs CD11c^WT^ (n = 4-7 mice per group).

**Figure S8.**
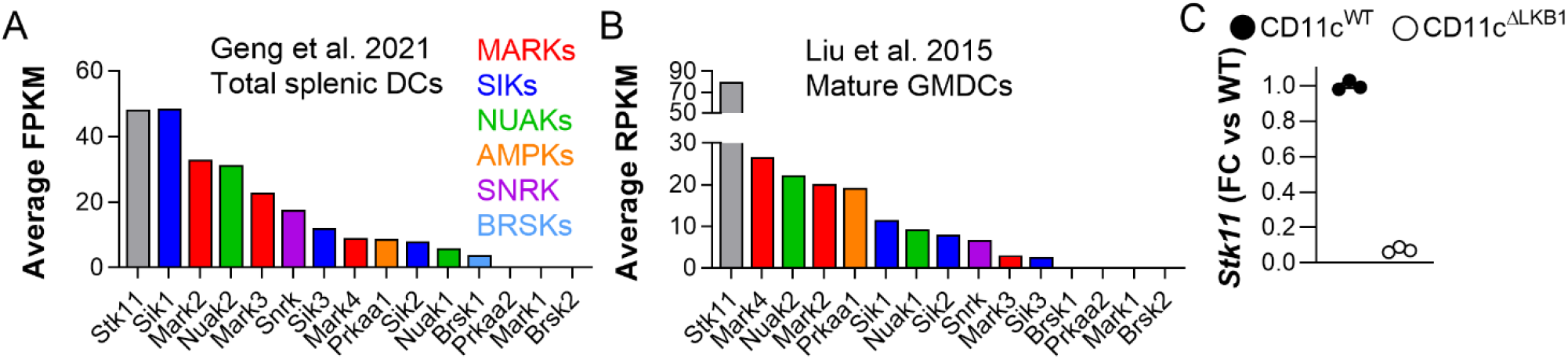
Transcriptional analysis of LKB1 and its substrates in DCs. **A-B:** Expression of *Stk11* (encoding LKB1) and its downstream targets *Mark1-4, Sik1-3, Nuak1-2, Prkaa1-2* (encoding AMPKα1-2), *Snrk* and *Brsk1-2* in total splenic DCs (Geng et al., *Immunology*. 2021; *A*) and mature GM-CSF-elicited bone marrow DCs (GMDCs; Liu et al., *J. Immunol*. 2015; *B*). **C:** *Stk11* expression in GMDCs from CD11c^WT^ (black symbols) and CD11c^ΔLKB1^ mice (open symbols) at day 10 of culture. Related to figure 5.

